# Diabetes Might Shape Vasculature, Tumor Dissemination and Circulating Inflammation Marker Profiles in Prostate Cancer

**DOI:** 10.64898/2026.04.27.720986

**Authors:** Kamil Kowalski, Marta Popęda, Julia Richert, Robert Wenta, Jolanta Szade, Anna Żaczek, Tomasz Kryczka, Mikołaj Frankiewicz, Kevin Miszewski, Marcin Matuszewski, Natalia Bednarz-Knoll

## Abstract

**Background and objective:** Chronic comorbidities like diabetes mellitus can influence cancer incidence and progression. However, their impact on pathological and molecular tumor characteristics, and dissemination, especially in prostate cancer (PCa), is not fully understood. Here, we investigated differences in the PCa molecular landscape with coexisting diabetes type 2 and its link to tumor dissemination.

**Methods:** D’Amico intermediate- or high-risk PCa patients (n=145), with type 2 diabetes or no diabetes, were analyzed for clinico-pathological features, circulating tumor cell (CTC) presence, yields and phenotypes in tumor-draining vein (TDVB) and peripheral blood (PB), primary tumor characteristics, and selected plasma biomarkers.

**Key findings and limitations:** Diabetes type 2 was found in 20/13.8% patients and associated with more advanced clinical outcomes, i.e. pathological tumor stage (p=0.011) and lymph node involvement (p=0.020). Among patients diagnosed before age 65, diabetes and PCa showed a borderline association with shorter time-to-biochemical recurrence (p=0.032). Diabetic PCa patients had significantly higher total CTC counts in TDVB but not in PB, indicating enhanced tumor cell dissemination from primary tumor (PT). PTs from diabetic PCa patients displayed a borderline association with higher ALDH1 expression (p=0.054) and significantly lower vascular vessel density (p=0.008). Similarly, in patients under 65, PTs from diabetic PCa patients expressed genes linked to decreased angiogenesis. Plasma analyses revealed elevated GDF15 levels in PB (p<0.001), increased TRAP5 concentrations in TDVB (p=0.001), and reduced osteonectin levels in TDVB (p=0.026) in diabetic PCa patients. The study’s limitation is the relatively small cohort, especially those with coexisting diabetes type 2.

**Conclusion and clinical implications:** Diabetes in PCa patients associates with advanced tumor stage, enhanced tumor cell dissemination, impaired vascularization, and distinct circulating biomarker alterations. Vascular alterations from diabetes, with systemic factors, especially in PCa patients under 65, may increase tumor dissemination in PCa. However, exact mechanisms need investigation in larger cohorts of patients.

**Graphical abstract:** 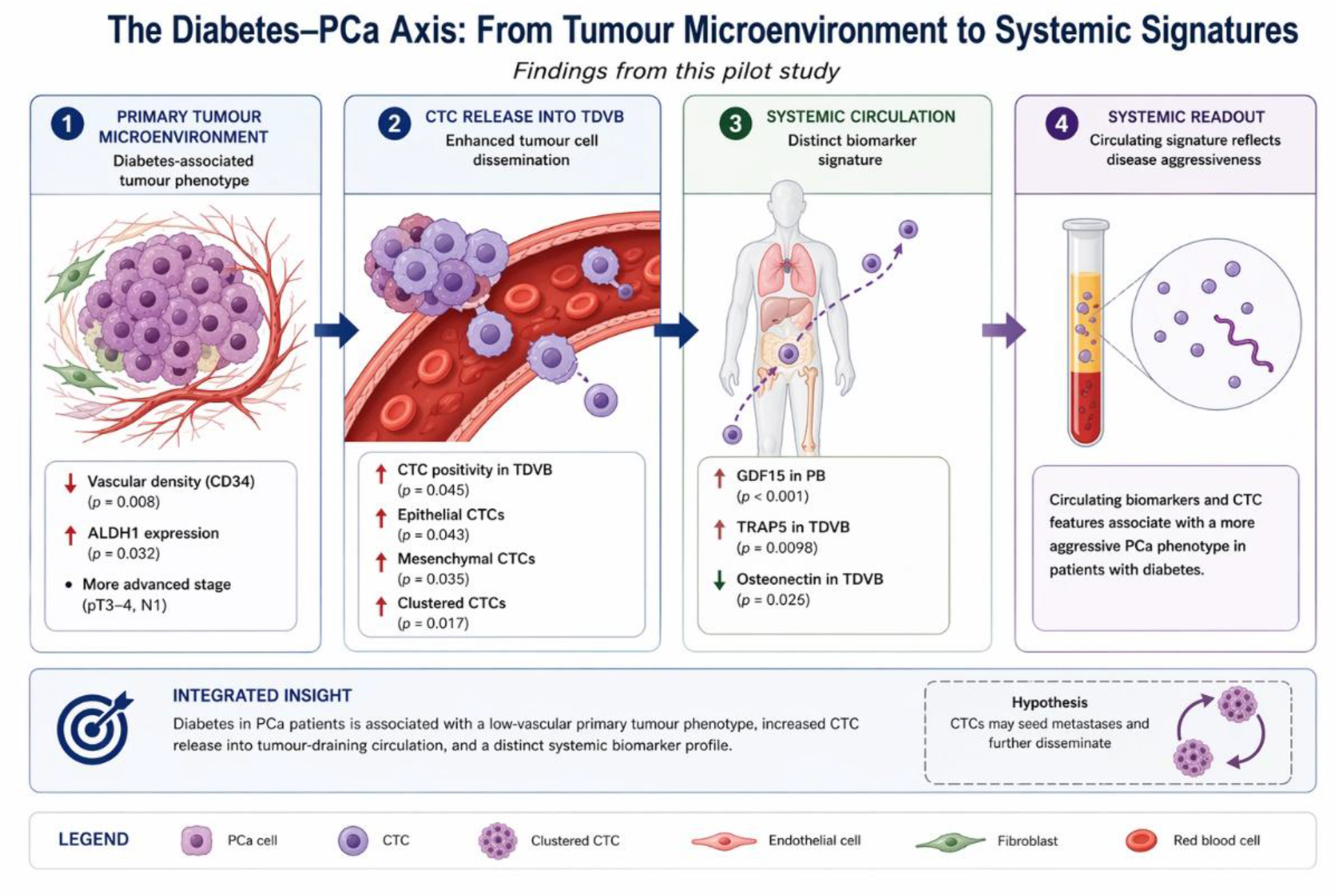

## Introduction

Chronic comorbidities are highly prevalent among tumor patients, including men diagnosed with PCa, particularly within older demographics. Of these comorbidities, diabetes mellitus warrants specific attention due to its intricate relationship with the incidence and progression of PCa. Epidemiological studies indicate that diabetes is associated with a decreased risk of developing PCa^1,2^. In contrast, once the disease manifests, diabetes may contribute to more aggressive tumor characteristics and poorer prognoses. This paradox is believed to arise from a combination of hormonal, metabolic, and inflammatory alterations associated with diabetes, including modified insulin and androgen signaling, chronic low-grade inflammation, and microvascular dysfunction^3,4^, mechanisms which might have a dual effect on tumor development. Nevertheless, the crosstalk between tumor development and different comorbidities remain inadequately understood^5^, and merits deepen investigation.

Previous research has predominantly concentrated on clinical outcomes and survival, with limited exploration of the underlying tumor biology and its dissemination processes in diabetic PCa patients. Specifically, there is a paucity of knowledge regarding the influence of diabetes on tumor dissemination and circulating blood-derived biomarkers, and even the molecular or histopathological characteristics of primary tumors. Given the growing interest in liquid biopsy methodologies and the significance of the tumor microenvironment in cancer progression, elucidating these associations holds both biological and clinical importance. Chronic hyperglycemia and diabetic microangiopathy are well-established drivers of vascular remodeling in other organs. We hypothesized that coexisting diabetes may reshape the prostatic tumor microenvironment and, through vascular and inflammatory alterations, influence the dissemination of tumor cells into the circulation

In this study, we investigated a cohort of d’Amico intermediate- and high-risk prostate cancer (PCa) patients to assess the potential influence of coexisting diabetes on tumor characteristics, dissemination, and biomarker profiles. Specifically, we compared clinico-pathological features, circulating tumor cells (CTCs) counts and phenotypes in peripheral blood (PB) and tumor-draining venous blood (TDVB)^6^, histological and molecular parameters of primary tumors, and circulating levels of selected biomarkers. By integrating tissue-based and liquid biopsy data, our objective was to provide a more comprehensive understanding of how diabetes may shape PCa progression.

## Materials and Methods

### Patients cohort

D’Amico intermediate and high-risk prostate cancer patients (EAU classification 2026) undergoing radical prostatectomy were recruited in the years 2018-2023 in the Department of Urology, University Clinical Center, Medical University of Gdańsk based on the informed written consent approved by the local Ethic Comitee (#NKBBN/286/2018, #NKBBN/748/2019–2020, and #NKBBN/ 434/2017) (Table 1). 20 patients (13.8%) had ongoing diabetes type 2, treated with metformin (500-2000mg) in 4 patients combined with other drugs (i.e. Dapagliflozin, Glimepiride, Empagliflozin, Gliclazide). The average duration of diabetes treatment among patients was 9.3 years, with a range spanning from 2 to 24 years. At hospital admission, non-fasting blood glucose level were measured as part of routine clinical assessment. Due to inconsistent availability, HbA1c values were excluded from the analysis. Primary tumors were collected and prepared as tissue microarrays as described^7^ for further immunohistochemical and multigene expression analysis, while matched peripheral blood samples were collected for CTC analysis and cytokine measurements.

**Table 1.** Patient demographics and clinical characteristics across the groups. Please, note that not all parameters sum up to 145 due to the missing data. PCa indicates prostate cancer, RP - radical prostatectomy, yr – years, BMI – body mass index, PSA – prostate specific antigen. Mann-Whitney or Chi2 test were used.

| Parameter | PCa without diabetes | PCa + diabetes | p-value |
| --- | --- | --- | --- |
| Patients, n (%) | 125 (86.2%) | 20 (13.8%) | - |
| Median age at RP (yr) | 66 (47-81) | 69 (51-77) | 0.375 |
| Patient under 65 years old (<65), n, % | 51 (40.8%) | 7 (35%) | - |
| BMI | 27.8 (18-41) | 29.7 (23-38) | 0.011 |
| Mean preoperative PSA (ng/ml) | 9.5 (2.2-43.36) | 11,8 (4,6-42) | 0.144 |
| Hypertension, n, % | 68 (54.4%) | 17 (85%) | 0.038 |
| Hypercholesterolemia, n, % | 19 (15.2%) | 5 (25%) | <0.001 |
| Cardiovascular disease, n, % | 27 (21.6%) | 7 (35%) | <0.001 |
| Thyroid disorders, n, % | 7 (5.6%) | 3 (15%) | <0.001 |

**Table 2.** Differentially expressed genes in prostate cancer patients.

| Gene | Non-diabetic<br>& <65 years | Diabetic<br>& <65 years | Log2FC | Classification | p-value |
| --- | --- | --- | --- | --- | --- |
| CDH11 | 157 | 410 | 1.38 | upregulated | 0.002 |
| SERPINF1 | 119 | 908 | 2.92 | upregulated | 0.006 |
| FBLN5 | 96 | 171 | 0.83 | upregulated | 0.011 |
| DENR | 120 | 262 | 1.13 | upregulated | 0.014 |
| CDS1 | 85 | 200 | 1.23 | upregulated | 0.017 |
| UBA52 | 688 | 1280 | 0.89 | upregulated | 0.020 |
| KDM1A | 80 | 157 | 0.96 | upregulated | 0.020 |
| ISLR | 74 | 200 | 1.42 | upregulated | 0.021 |
| TFDP1 | 117 | 194 | 0.73 | upregulated | 0.021 |
| TIMP2 | 178 | 449 | 1.33 | upregulated | 0.027 |
| DCN | 230 | 698 | 1.60 | upregulated | 0.027 |
| VHL | 210 | 275 | 0.39 | upregulated | 0.027 |
| BGN | 514 | 1341 | 1.38 | upregulated | 0.027 |
| PRR15L | 56 | 101 | 0.85 | upregulated | 0.030 |
| FGFR2 | 62 | 110 | 0.83 | upregulated | 0.033 |
| NR3C1 | 161 | 342 | 1.08 | upregulated | 0.033 |
| ARHGDIB | 452 | 927 | 1.03 | upregulated | 0.046 |
| DPT | 51 | 117 | 1.18 | upregulated | 0.049 |
| PMP22 | 51 | 127 | 1.31 | upregulated | 0.049 |
| PTPRM | 57 | 97 | 0.77 | upregulated | 0.049 |
| SPARC | 217 | 553 | 1.35 | upregulated | 0.049 |

| Gene | ≥65 years<br>& diabetic | <65 years<br>& diabetic | Log2FC | Classification | p-value |
| --- | --- | --- | --- | --- | --- |
| DENR | 117 | 262 | 1.16 | upregulated | 0.017 |
| UBA52 | 625 | 1280 | 1.03 | upregulated | 0.017 |
| KDM1A | 75 | 157 | 1.06 | upregulated | 0.017 |
| TIMP2 | 182 | 449 | 1.30 | upregulated | 0.017 |
| FGFR2 | 59 | 110 | 0.89 | upregulated | 0.017 |
| NR3C1 | 112 | 342 | 1.60 | upregulated | 0.017 |
| AKT2 | 176 | 470 | 1.41 | upregulated | 0.017 |
| CDK14 | 127 | 811 | 2.67 | upregulated | 0.017 |
| TNS1 | 157 | 365 | 1.21 | upregulated | 0.017 |
| COL6A1 | 104 | 328 | 1.65 | upregulated | 0.017 |
| CDKN1A | 120 | 566 | 2.23 | upregulated | 0.017 |
| AKAP12 | 52 | 99 | 0.92 | upregulated | 0.017 |
| NME4 | 267 | 1945 | 2.86 | upregulated | 0.017 |
| ILK | 60 | 171 | 1.50 | upregulated | 0.017 |
| CLIC4 | 194 | 314 | 0.69 | upregulated | 0.017 |
| FSTL1 | 481 | 1552 | 1.69 | upregulated | 0.017 |
| DDR2 | 74 | 225 | 1.59 | upregulated | 0.017 |
| ALDOA | 600 | 2261 | 1.91 | upregulated | 0.017 |
| SMURF1 | 44 | 105 | 1.24 | upregulated | 0.017 |
| ID1 | 53 | 107 | 1.00 | upregulated | 0.017 |
| PTGS2 | 45 | 126 | 1.47 | upregulated | 0.017 |
| ISLR | 80 | 200 | 1.31 | upregulated | 0.022 |
| KRAS | 88 | 47 | -0.89 | downregulated | 0.022 |
| SRF | 99 | 229 | 1.20 | upregulated | 0.022 |
| PTRF | 18 | 208 | 3.46 | upregulated | 0.022 |
| PTK2 | 54 | 132 | 1.27 | upregulated | 0.022 |
| CHP1 | 64 | 115 | 0.84 | upregulated | 0.022 |
| CDH11 | 170 | 410 | 1.27 | upregulated | 0.033 |
| SERPINF1 | 115 | 908 | 2.97 | upregulated | 0.033 |
| CDS1 | 65 | 200 | 1.61 | upregulated | 0.033 |
| ARHGDIB | 214 | 927 | 2.11 | upregulated | 0.033 |
| DPT | 33 | 117 | 1.80 | upregulated | 0.033 |
| GIMAP4 | 42 | 99 | 1.22 | upregulated | 0.033 |
| ENO1 | 366 | 1797 | 2.29 | upregulated | 0.033 |
| ITGA8 | 50 | 91 | 0.85 | upregulated | 0.033 |
| PLEKHO1 | 86 | 169 | 0.97 | upregulated | 0.033 |
| MAF | 35 | 90 | 1.34 | upregulated | 0.033 |
| PCOLCE | 38 | 198 | 2.35 | upregulated | 0.040 |
| WIPF1 | 35 | 88 | 1.31 | upregulated | 0.040 |
| DICER1 | 119 | 75 | -0.66 | downregulated | 0.040 |

### Blood collection and processing

TDVB was obtained intraoperatively during radical prostatectomy, prior to prostate excision, from veins draining the tumor-involved prostate lobe, whereas peripheral blood (PB) was collected preoperatively CTCs were isolated, enumerated and phenotyped as described^8^. Briefly, EDTA-blood samples (vol. of up to 7.5 mL) were collected from all recruited patients and processed as soon as possible after blood donation (with 68% of the samples processed within up to 3 h after donation). Initially, 2-3 mL of blood was discarded to prevent contamination by skin cells (such as keratinocytes and fibroblasts) and endothelial cells during venipuncture. The (peripheral) blood mononuclear cell ((P)BMC) fraction was isolated through density gradient centrifugation. Briefly, the blood was diluted with 1x phosphate-buffered saline (1xPBS) to a total volume of 6 or 12 mL, layered onto Ficoll® 400 (Sigma-Aldrich, St. Louis, MO, USA), and centrifuged at 400× g for 30 minutes with the brake off. The (P)BMC fraction was then collected, fixed in 4% formaldehyde, and stored at −80°C in aliquots for subsequent CTC analysis, while matched plasma samples - at -80°C in aliquots for subsequent cytokines measurements.

### CTC isolation and analysis

CTC isolation and analysis was performed as described^8^. The (P)BMCs under investigation were thawed and subsequently washed with 1 mL of 1x PBS to eliminate formaldehyde. They were then incubated for 30 minutes at +4 °C with an antibody cocktail. Immunofluorescent staining of (P)BMCs fractions was conducted utilizing a combination of antibodies to exclude normal cells (based on 4 markers, i.e. CD45, CD31, aSMA and CD29), while identify and phenotype putative circulating tumor cell (CTC): pan-keratins (K, AE1/AE3 clone, AF488-conjugated, Thermo Fisher Scientific, Waltham, MA, USA, #53-9003-82; C11 clone, AF488-conjugated, Thermo Fisher Scientific, #MA5-18156) and vimentin (V, D21H3 clone, AF647-conjugated, Cell Signaling, Danvers, MA, USA, #9856); CD29 (TS2/16 clone, SuperBright600-conjugated, Thermo Fisher Scientific, #63-0299-42); CD45 (REA747 clone, APC-Vio770-conjugated, Miltenyi Biotec, North Rhine-Westphalia, Germany, #130-110-635); and CD31 (WM59 clone, APC-Cy7-conjugated, BioLegend, San Diego, CA, USA, #303120). These antibodies were diluted at ratios of 1:2500, 1:2500, 1:50, 1:100, 1:1000, 1:50, and 1:10, respectively, in 1x Perm-Wash Buffer (BD Biosciences, Franklin Lakes, NJ, USA).

Following an additional washing step in 1x PBS, the cells were resuspended in 30 μL of 1x PBS, counterstained with DAPI (BD Biosciences, 1 μg/mL), and immediately subjected to analysis using imaging flow cytometry. Imaging flow cytometry analysis was conducted using the Amnis® ImageStream® X Mk II (Cytek Biosciences), which is equipped with lasers at 405 nm, 488 nm, and 642 nm, and utilizes INSPIRE™ software (version 200.1.681.0; Cytek Biosciences) for data acquisition. The subsequent analysis of the acquired data was performed using IDEAS software (version 6.2; Cytek Biosciences). The cells were imaged at 40× magnification at a low speed to ensure high-quality images. For markers potentially coexisting within the same subpopulation of cells and detected using the same laser (i.e., V, CD45, CD31), and for highly intense fluorescence in the DAPI channel, optimal compensation was applied.

The optimized parameter settings for acquisition were consistently applied to all analyzed samples. Each analyzed cell was visually verified in every fluorescent channel based on fluorescence intensity and pattern to exclude autofluorescence and/or fluorescence crosstalk. Only distinct fluorescence patterns against the background were considered true events. All DAPI+ cells were counted and served as the initial gate, while DAPI+/CD45−CD31−/aSMA-/CD29-cells were considered as putative CTCs and further phenotyped. To gate potential CTCs, the fluorescence intensities of K and V were visualized in a two-dimensional dot plot. The potential preselected CTCs were visually confirmed from immunofluorescence images for their morphology and staining details, tagged, and manually counted. The detected cells were classified as follows: epithelial CTCs if K+/V−, mesenchymal CTCs if K−/V+, epithelial–mesenchymal CTCs if K+/V+, and negative for both epithelial and mesenchymal markers if K−/V−.

### Cytokine/Mediators analysis

The concentrations of selected plasma cytokines (i.e. DKK-1, GDF-15, OPG, osteonectin/SPARC, periostin, and TRAP5) were quantified in collected plasma from PB and TDVB samples using a commercially available magnetic microbead–based 6-plex assay (MILLIPLEX MAP Human Cancer/Metastasis Biomarker Magnetic Bead Panel, Merck). Measurements were performed on a Bio-Plex® 200 system utilizing Luminex xMAP technology (Bio-Rad) in accordance with the manufacturer’s instructions, with plasma diluted 1:4. Plasma samples were analyzed in a blinded manner. Cytokine/mediator concentrations were calculated based on seven-point standard curves with descending concentrations, within an experimental range of 0.01–11,467 ng/mL, depending on the cytokine/mediator.

### Immunohistochemistry staining for vascular vessel numbers assessment

Initially, tissue microarray (TMA) sections, with a thickness of 4 to 6 μm, were deparaffinized, followed by antigen retrieval using the EnVision FLEX Target Retrieval Solution Low pH (Dako Agilent) in a PT Link device (PT200, Dako Agilent) for 20 min. at 97°C, then incubated in Peroxidase-Blocking Solution (Dako) for 5 min. The sections were incubated for 1 h at RT with ready-to-use mouse monoclonal anti-CD34 antibody (clone QBEnd10, Dako) envisioned by EnVision Kit, Rabbit/Mouse (Dako) and counterstained with haematoxylin (Sigma Aldrich). Staining was evaluated by quantifying the number of CD34-positive vascular vessels with a visible lumen in the investigated tumor area, as described ^9^ . Immunohistochemical staining was assessed at 200× magnification using a light microscope (Olympus BX 43, Olympus, Japan). The total number of CD34-positive vessels with a visible lumen was recorded for each tumor sample. The obtained values were categorized based on mathematical cut-offs (quartiles) as follows: q4 as low, q3,2 as - medium, and q1 as - high number of vascular vessels within investigated area of a tumor.

### Immunofluorescence staining

First, 4-to 6-μm-thick TMA sections were deparaffinized, and antigen retrieval was performed using EnVision FLEX Target Retrieval Solution Low pH (Dako Agilent) in a PT Link device (PT200, Dako Agilent) for 20 min at 97°C. The TMAs were then incubated overnight at 4°C with a rabbit anti-ALDH1 antibody (Abcam, ab52492) diluted 1:100 prior to incubation with a secondary antibody for 60 min. Nuclei were stained with DAPI, and the sections were mounted with Vectashield Antifade Mounting Medium (Vector Laboratories). ALDH1 expression was evaluated in tumor cells within the investigated tumor area based on intensity (none, weak, moderate, strong) and the percentage of total tumor cells.

### NanoString nCounter gene expression assay

RNA isolation and profiling using NanoString technology was performed as described ^7^. Briefly, total RNA was extracted from 69 samples of formalin-fixed paraffin-embedded (FFPE) TMA-originated tissue cores from primary prostate tumors, preselected for TMA by the experienced pathologist, consisting of three 20 μm-thick, unstained FFPE sections per patient, utilizing a RNeasy Mini Kit (QIAGEN, Hilden, Germany) in accordance with the manufacturer’s protocol. The integrity of the RNA was evaluated using an Agilent 2100 Bioanalyzer (Agilent Technologies, Santa Clara, CA, USA) with an Agilent RNA 6000 Pico Kit (Agilent Technologies). A volume of 4 μL of the extracted RNA was preamplified using the nCounter Low RNA Input Kit (NanoString Technologies, Seattle, WA, USA) with a specific primer pool targeting the sequences of 740 genes included in the nCounter Cancer Progression Profiling Panel (NanoString Technologies). The preamplified samples were subsequently analyzed using the NanoString nCounter Analysis System (NanoString Technologies) following the manufacturer’s protocols for hybridization, detection, and scanning. Background correction and data normalization were conducted using nSolver 4.0 software (NanoString Technologies). The background level was determined by setting a threshold above the mean plus two standard deviations of negative control counts, and the data were normalized based on the global mean of the counts of the positive control probes included in the assay and the four most stably expressed housekeeping genes, specifically CNOT4, HDAC3, DDX50, and CC2D1B. Differentially expressed genes (DEGs) were identified using a p < 0.05 threshold (Mann–Whitney–Wilcoxon test), and median-based log2FC values were reported. Functional annotation of DEGs was performed using GO BP (Gene Ontology Biological Process) terms via the STRING database^10^.

### Statistical analysis

Statistical analyses were conducted using the SPSS 27.0.1.0 software package (SPSS Inc., Chicago, IL, USA), licensed to the University of Gdańsk. The precise counts of circulating tumor cells (CTCs) were calculated per 1 million (P)BMCs to ensure sample normalization. Differences in CTC enumeration among cancer patients with varying clinicopathological parameters were assessed using Mann-Whitney and Kruskal-Wallis tests. Survival analysis was performed using GraphPad Prism and visualised using Kaplan-Meier curves. All statistical analyses were two-sided, with a p-value of less than 0.05 considered statistically significant.

## Results

At least one of the documented chronic diseases including diabetes, cardiological diseases (i.e. atrial fibrillation, hypertension), hypercholesteremia, or thyroid disorder was observed in 64 of 145 (44.1%) recruited d’Amico intermediate or high risk PCa patients. In our study, diabetes but not other comorbidities, was associated with more advanced characteristics of PCa. Diabetes is known to lower the odds of PCa occurrence but on the other side also to worsen its course^2^. Therefore, here, we explored the potential impact of coexisting diabetes on clinico-pathological features of a tumor, tumor dissemination, and selected molecular markers investigated within primary tumors or in plasma derived from PB and TDVB of those patients.

Diabetes was documented in 20 (13.8%) PCa patients. Patients with coexisting PCa and diabetes demonstrated more advanced pathological tumor stages, evidenced by a higher frequency of tumors characterized as pT3 and pT4 (p=0.011, **Fig. 1A**) and more frequent lymph node involvement (N1 status, Fisher’s exact test, p=0.020, **Fig. 1B**). Interestingly, despite these indicators of advanced disease, diabetic PCa tumors were mainly characterized by Gleason score 4+3 in contrast to 3+4 tumors and even those with >7 (p=0.026, **Fig. 1C**). Diabetes seemed to co-occur more frequently in PCa patients diagnosed at the older age (p=0.042, **Fig. 1D**). However, only in patients diagnosed under 65 years of age, we observed a borderline association with the shorter time to biochemical recurrence (BCR) when PCa and diabetes were coexisting (p= 0.0215, **Fig. 1E, F**).

**Fig. 1.**
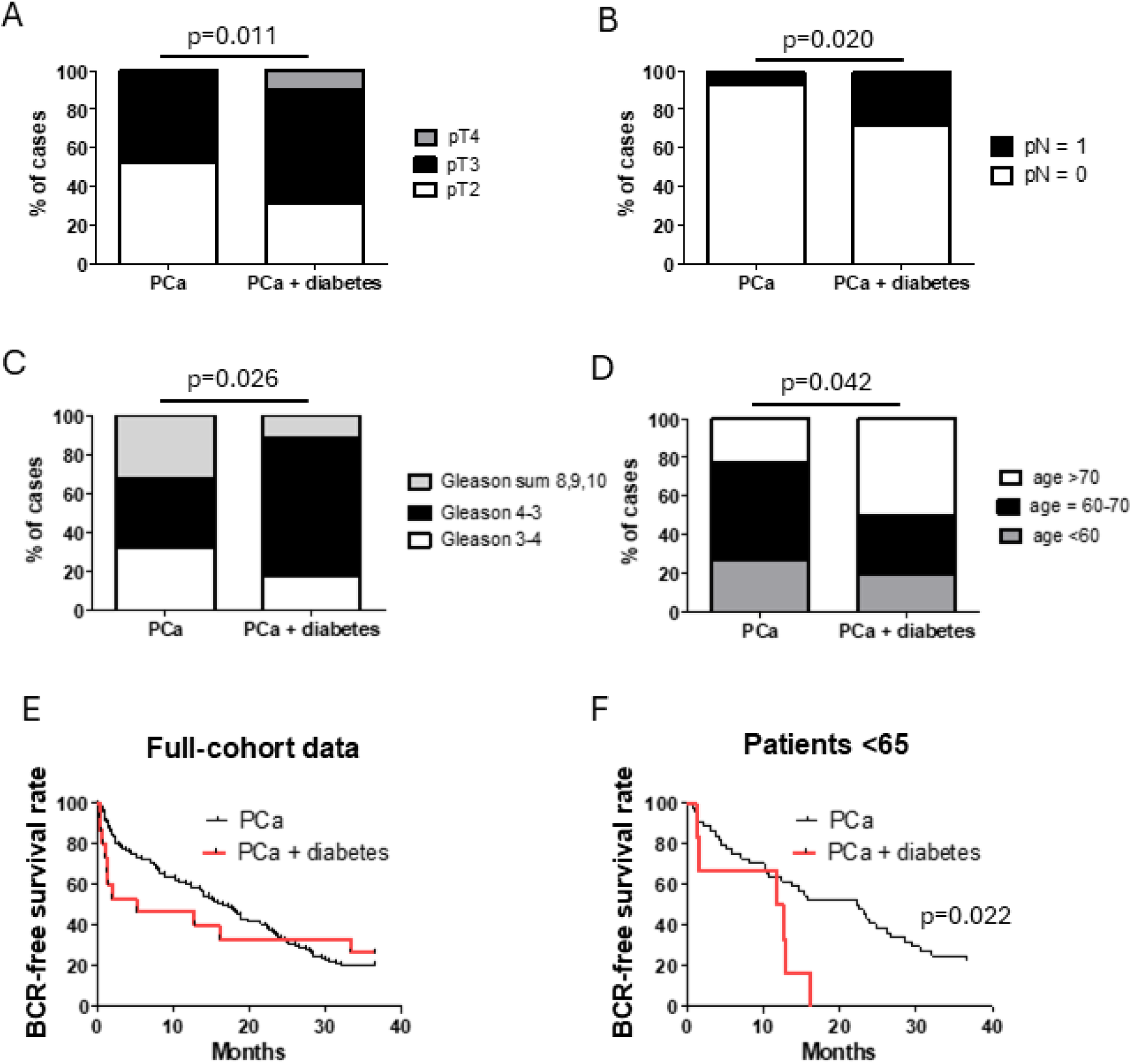
Clinico-pathological parameters of diabetic PCa patients. (A) Proportion of tumor stages in patients with prostate cancer (PCa, n=119) and those with both PCa and diabetes (n=19); (B) Proportion of lymph node involvement status in patients with PCa (n=118) and those with both PCa and diabetes (n=18); (C) Proportion of Gleason scores in patients with PCa (n=118) and those with both PCa and diabetes (n=17); (D) Distribution of age groups among patients with PCa (n=124) and those with both PCa and diabetes (n=18); (E) Time to biochemical recurrence in patients with PCa (n=109) and those with both PCa and diabetes (n=17); (F) Time to biochemical recurrence in patients below 65 years old with PCa (n=10) and those with both PCa and diabetes (n=5).

Upon evaluating the status of tumor dissemination, it was observed that a greater proportion of diabetic PCa patients exhibited CTCs in TDVB (p=0.045), but not in PB, compared to PCa patients without diabetes (**Fig. 2A-B, Suppl. 1. A-B**). This was also true for individually counted epithelial (K+V-) CTCs (p=0.043), putative EMT-like (in particular, mesenchymal, i.e. K-V+, p=0,035, and negative, (K-V-) CTCs, p=0.033), and clustered CTCs in TDVB (p=0.017), but not in PB (**Fig. 2C, Suppl. 1. C**). The higher percentage of diabetic patients exhibiting CTCs in the bloodstream, particularly in TDVB, suggests an intensified dissemination of tumor cells from the primary tumor.

**Fig. 2.**
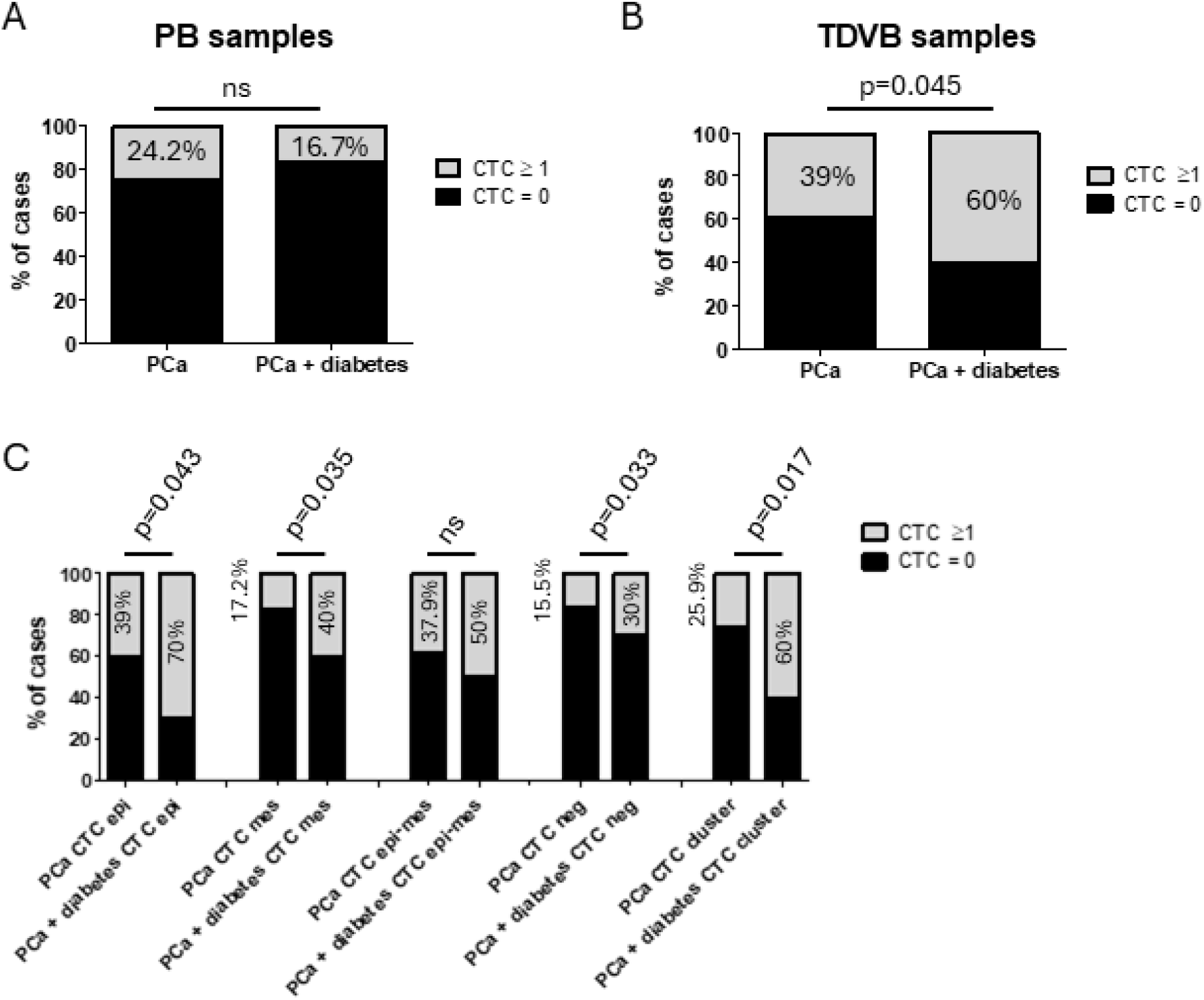
CTC analysis in PCa patients. (A) Proportion of patients with prostate cancer (PCa, n=95) and those with both PCa and diabetes (n=18) exhibiting at least one circulating tumor cell (CTC) detected in peripheral blood (PB); (B) Proportion of patients with PCa (n= 59) and those with both PCa and diabetes (n=10) with at least one CTC detected in blood from the tumor-draining vein (TDVB); (C) Proportion of patients with PCa (n=58) and those with both PCa and diabetes (n=10) exhibit at least one epithelial CTC, one epithelial-mesenchymal transition (EMT)-like CTC, one negative CTC, or one clustered CTC in TDVB.

Selected plasma biomarker measurement in PB and TDVB of patients indicated elevated levels of growth differentiation factor 15 (GDF15) in PB samples of diabetic patients (p<0.001, **Fig. 3A**) and higher concentrations of tartrate-resistant acid phosphatase 5 (TRAP5) in TDVB (p=0.0098, **Fig. 3B**). Conversely, plasma osteonectin levels were lower in TDVB of diabetic individuals (p=0.026, **Fig. 3C**). No correlation was observed between diabetes and the levels of DKK-1, OPG, and periostin in either PB or TDVB.

**Fig. 3.**
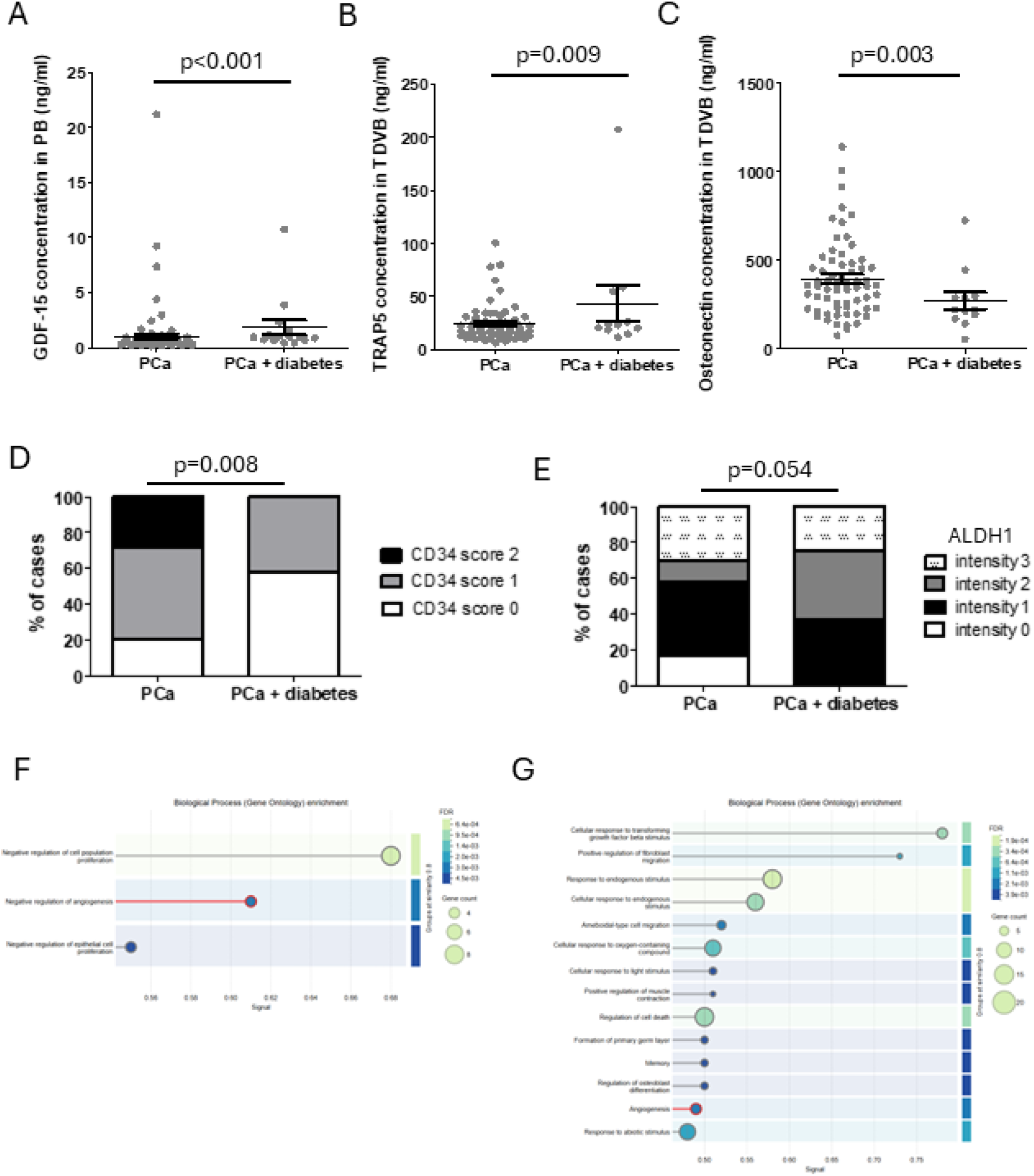
Plasma biomarkers in PB and TDVB, and molecular profiels of primary tumors in PCa patients. (A) Concentration of GDF-15 in peripheral blood among patients with PCa (n=115) and those with both PCa and diabetes (n=15); (B) Concentration of TRAP5 in blood from the tumor-draining vein among patients with PCa (n=65) and those with both PCa and diabetes (n=11); (C) Concentration of osteonectin in blood from the tumor-draining vein among patients with PCa (n= 64) and those with both PCa and diabetes (n=11); (D) Proportion of patients with PCa (n=56) and those with both PCa and diabetes (n=10), categorized by number of vascular vessels identified using CD34 staining. (E) Proportion of patients with PCa (n=86) and those with both PCa and diabetes (n=8), categorized by ALDH1 protein levels as none, weak, medium or strong; (F) Gene ontology (GO) analyses of the NanoString nCounter gene expression assay in samples from PCa patients under 65 years of age, comparing those with diabetes (n=3) to those without diabetes (n=22) visualised using STRING database; (G) Gene ontology (GO) analyses of the NanoString nCounter gene expression assay in samples from PCa patients with diabetes, comparing those under 65 years of age (n=3) to those over 65 years old (n=7) visualised using STRING database.

Primary tumors of diabetic PCa patients showed a significantly lower number of vascular vessels compared to non-diabetic counterparts (p=0.008, **Fig. 3D**) and association with more frequently occurring higher expression of ALDH1 (p= 0.032, **Fig. 3E**) but not CD44+/CD24-phenotype (data not shown). Multigene transcriptomic profiling of primary PCa tissue fragments using NanoString technology did not reveal significant differences between patients with and without diabetes (data not shown).

However, as shorter time to BCR was observed to be associated with co-existence of diabetes exclusively in younger patients (<65), we focused our further analysis on comparison of patients stratified simultaneously according to age and diabetes. In PCa patients under 65 years of age, comparison of diabetic and non-diabetic cases identified 21 upregulated genes. These were associated with processes such as cell proliferation and notably included four genes involved in the negative regulation of angiogenesis (i.e., DCN, SPARC, SERPINF1, PTPRM; **Fig. 3F, Suppl. Tab. 2**), suggesting a shift toward anti-angiogenic signalling in diabetic patients. Within the diabetic subgroup, comparison of younger and older patients revealed 38 upregulated genes in younger individuals. These were associated, among others with cell migration, and included six genes linked to angiogenesis (i.e., FGFR2, PTK2, PTGS2, CLIC4, SRF, ID1; **Fig. 3G, Suppl. Tab. 2**), indicating relatively increased angiogenic activity in younger diabetic PCa patients.

## Discussion

Risk stratification is essential in the management of prostate cancer, as radical treatment should primarily be considered for patients at high risk of metastasis and rapid disease progression. Decisions should be made by factors such as life expectancy, disease risk category, and overall clinical condition. Incorporating comorbidities, which frequently coexist with age-associated malignancies, as confounding factors in treatment decision-making requires a comprehensive understanding of the interactions between these disorders and tumors. Diabetes, as one of the most significant comorbidities affecting overall metabolism, can serve as a major predictive factor for tumor behavior, and was shown to modulate tumor development and even progression in many correlative studies. To the best of our knowledge, this is the first study dissecting the broader molecular landscape of PCa when coexisting with diabetes and showing decreased vasculature while increased seeding in diabetic PCa patients.

While diabetes co-occurred only in 14% of PCa cases, it was associated with advanced disease characteristics, distinct vasculature-related tumor profiles, and increased tumor cell dissemination via blood circulation. In alignment with prior studies^11^, our research found that diabetic PCa patients were diagnosed at an older age and had more aggressive disease, with higher pT3-4 tumors and lymph node involvement. Despite advanced stage, primary tumors in diabetic PCa patients showed predominantly Gleason score 4+3, indicating a distinct pattern even in comparison to patients with Gleason score >7. This aligns with research showing diabetes decreases PCa risk but may worsen progression and may be explained by altered hormonal and metabolic conditions promoting tumor aggressiveness while influencing differentiation^12^.

Next, we analyzed PCa dissemination in the context of diabetes. The ability to analyze not only PB but also TDVB is unique and with greater precision reveals seeding from primary tumors, allowing us to demonstrate for the first time that diabetic PCa patients may be indeed characterized by increased tumor cell dissemination. We found elevated CTC counts, including epithelial, EMT-like, and clustered CTCs in TDVB, which suggests enhanced tumor spread in diabetic patients. The increased CTC dissemination may reflect a permissive microenvironment with metabolic and inflammatory changes facilitating tumor cells survival. Indeed, analysis of primary tumors showed decreased vascular density in diabetic PCa, possibly reflecting adaptation to hyperglycemia or vascular pathology commonly associated with diabetes mellitus, including microangiopathy. Such microenvironment with decreased density of vascular vessels can potentially generate hypoxia^9^ and other low vascularity-associated pathways, initiating or at least boosting the ability of tumor cells to migrate.

Taken together, the reduced microvessel density, the enrichment of mesenchymal and clustered CTCs in the tumor-draining vein, and the systemic elevation of GDF15 point toward a coherent mechanistic model in which diabetes-driven microangiopathy might generate focal hypoxia within the primary tumor, favor epithelial–mesenchymal plasticity, and, via systemic pro-inflammatory mediators, create conditions permissive to intravasation and early seeding. The fact that these changes were most pronounced in younger patients is consistent with the hypothesis that cumulative metabolic stress is not the sole driver; rather, the interaction between diabetes and tumor biology may be particularly consequential when it occurs on a younger vascular and immunological background.

Of note, in plasma biomarkers evaluation, diabetic PCa patients showed elevated growth differentiation factor 15 (GDF15) in PB and increased tartrate-resistant acid phosphatase 5 (TRAP5) in TDVB, while osteonectin levels were reduced. GDF15 influences both metabolic regulation and cancer progression, reflecting among others diabetic conditions and tumor responses. TRAP5 elevation may indicate early metastatic processes^13^, while reduced osteonectin could suggest altered stromal interactions in diabetic patients^14^. GDF15 is well known to be connected to diabetes and inflammation process^15^ and may exhibit a dual role in PCa, acting as a tumor suppressor in early stages by inhibiting proliferation and inducing apoptosis, while in advanced disease it promotes tumor progression and bone metastasis through activation of osteoclastogenesis and enhancing invasive pathways^16^. This dual function is reflected clinically, where higher GDF15 levels in primary tumors correlate with better prognosis, whereas reduced expression is associated with metastasis and poor outcomes^17,18^. GDF15 was increased in PB not in TDVB of diabetic PCa patients which might suggest the systemic diabetes-related background of the GDF15 secretion in those patients, rather than secretion of this mediator from primary tumors. However, it might be hypothesized that such a systemic diabetes-related pro-inflammatory mediator, might still influence the vasculature within tumor and even formation of premetastatic niche in those patients.

Transcriptomic profiling using NanoString technology found no significant gene expression differences between diabetic and non-diabetic tumors, suggesting that phenotypic variations may be, indeed, caused by microenvironmental factors rather than intrinsic molecular profile of tumor cells itself. On the other hand, when analyzing only patients under the age of 65, a group of patients in which diabetes was associated with shorter time-to-biochemical recurrence, we identified several dysregulated pathways based on transcriptomic profiling. One of the most pronounced processes was angiogenesis, which occurred to be downregulated in this cohort of patients (i.e. diabetic PCa patients under 65). This finding corresponds with the decreased vasculature identified in our study using immunohistochemistry and CD34 staining in diabetic PCa patients. This finding potentially highlights the necessity of considering diabetes status in the management of PCa patients and reducing their risk of progression.

A further consideration is the potential influence of antidiabetic pharmacotherapy. All diabetic patients in our cohort were receiving metformin, alone or in combination with SGLT2 inhibitors or sulfonylureas. Metformin has been repeatedly associated with modulation of PCa biology^19^, including effects on proliferation, angiogenesis, and the AMPK–mTOR axis, while SGLT2 inhibitors have more recently been implicated in metabolic reprogramming of the tumor microenvironment^20^. It is therefore conceivable that the tumor phenotype observed here represents not the effect of diabetes per se, but the combined signature of hyperglycemia and its treatment. Dedicated studies stratifying patients by drug class and treatment duration will be required to disentangle these contributions.

Our study acknowledges certain limitations. The sample size of patients with coexisting diabetes is relatively small, necessitating further research with larger cohorts from multiple centres. Additionally, future studies should encompass various types of diabetes, its treatments, and treatment duration before PCa diagnosis.

## Conclusions

Diabetes exerts a significant influence on the tumor microenvironment, contributing to a complex interplay that might affect the progression of PCa. Diabetes is well known to modulate the vascular architecture within tumors and, hence, might promote the intravasation of tumor cells into the bloodstream, thereby increasing the potential for metastasis. Given these mechanistic insights, it is advisable to incorporate diabetes status into risk stratification models for PCa management. Understanding the ways in which diabetes alters tumor biology could inform personalized therapeutic approaches. Our findings suggest that patients younger than 65 years could benefit from integrating diabetes considerations into PCa treatment decisions. Prospective studies incorporating glycemic control metrics, antidiabetic treatment data, and larger diabetic subgroups, particularly among patients under 65, are warranted to determine whether diabetes status should be formally integrated into existing PCa risk stratification models.

## List of abbreviations

ALDH1: aldehyde dehydrogenase 1
BCR: biochemical recurrence
CTC: circulating tumor cells
EMT: epithelial-mesenchymal transition
FFPE: formalin-fixed paraffin-embedded
GDF15: growth differentiation factor 15
IHC: immunohistochemistry
PB: peripheral blood
(P)BMCs: (peripheral) blood mononuclear cells
PBS: phosphate-buffered saline
PCa: prostate cancer
pT: pathological tumor stage denotes the size and extent of the primary tumor
PT: primary tumor
TDVB: tumor-draining vein blood
TMA: tissue microarrays
N status: lymph node involvement status
TRAP5: tartrate-resistant acid phosphatase 5

## Ethics approval and consent to participate

This study was conducted in accordance with the Declaration of Helsinki and approved by the Independent Bioethics Committee for Scientific Research at the Medical University of Gdansk (study protocols: #NKBBN/286/2018, #NKBBN/748/2019–2020, and #NKBBN/ 434/2017). The patients provided written informed consent to participate in the study.

## Consent for publication

Not applicable

## Availability of data and materials

Representative images of CTCs as well as linked gene expression data are available on the webpage: www.CTCatlas.org. The raw datasets used and/or analyzed during the current study are available from the corresponding author on reasonable request.

## Authors’ contributions

Study conceptualization, design and supervision: NBK; patients recruitment: MM, MF, KM; experiments and analysis: KK, MP, JR, RW, TK, NBK; data curation: MP, JR, RW, TK; resources: AZ, NBK; funding and administrative management: NBK; draft writing: KK, NBK. All authors read and approved the final manuscript.

## Acknowledgements

Th authors thank all patients who agreed to participate in the study, and Dr Beata Pieczyńska-Uziębło for her help with staining the blood samples.

