## Supplementary material for "Diabetes Might Shape Vasculature, Tumor Dissemination and Circulating Inflammation Marker Profiles in Prostate Cancer": Suplemental Figure 1

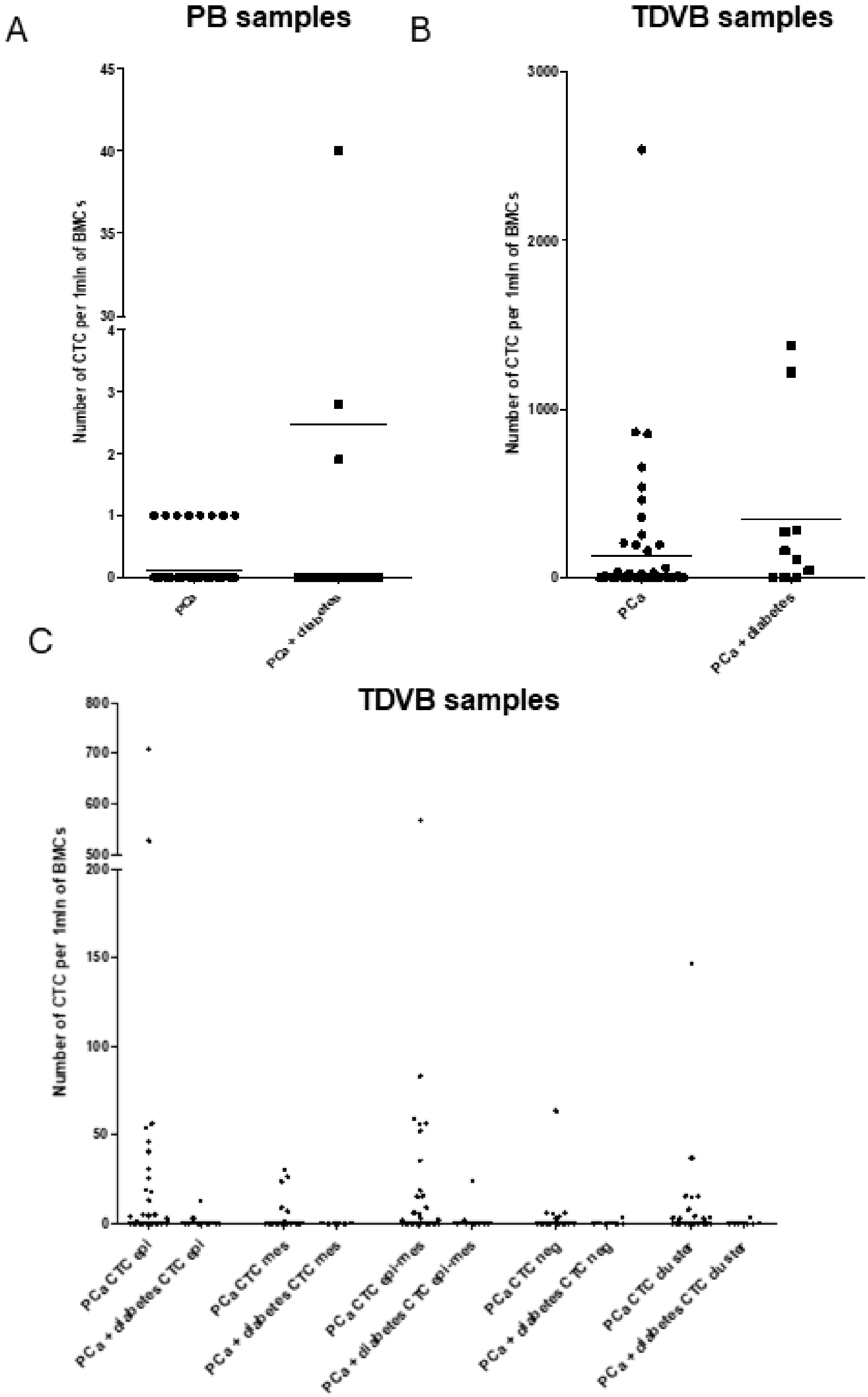
A
PB samples
Number of CTC per 1mln of BMCs
45
40
35
30
4
3
2
1
0
PCa
PCa + diabetes
B
TDVB samples
Number of CTC per 1mln of BMCs
3000
2000
1000
0
PCa
PCa + diabetes
C
TDVB samples
Number of CTC per 1mln of BMCs
800
700
600
500
200
150
100
50
0
PCa CTC epi
PCa + diabetes CTC epi
PCa CTC mes
PCa + diabetes CTC mes
PCa CTC epi-mes
PCa + diabetes CTC epi-mes
PCa CTC neg
PCa + diabetes CTC neg
PCa CTC cluster
PCa + diabetes CTC cluster

**Supplementary Figure 1. Number of CTCs in PCa patients** (A) The number of circulating tumor cells (CTCs) per one million peripheral blood mononuclear cells (PBMC) in patients with prostate cancer (PCa, n=95) and those with both PCa and diabetes (n=18) in peripheral blood (PB); (B) The number of CTCs per one million blood mononuclear cells (BMC) in patients with PCa (n=59) and those with both PCa and diabetes (n=10) in tumor-derived vascular blood (TDVB); (C) The number of CTCs per one million BMCs in patients with PCa (n=58) and those with both PCa and diabetes (n=10), exhibiting epithelial, epithelial-mesenchymal transition (EMT)-like CTC, negative CTC, or clustered phenotypes in TDVB.
